# RidgeTrace: extracting toothed whale whistle contours without annotated training data

**DOI:** 10.64898/2026.11.22.768378

**Authors:** Venkata Naga Balarama Murthy Pelluri, Sri Parjanya Bejjanki, Shaik Mohammed Navab Alisha, Shashank Sanga, Brindanjali Thumu, Nagarani Korla

**Affiliations:** Department of Computer Science and Engineering, Neil Gogte Institute of Technology, Hyderabad, Telangana, India

**Keywords:** Dolphin whistles, automatic detection, contour extraction, deep learning, synthetic training data, passive acoustic monitoring

## Abstract

Few toothed whale recordings have the whistle-by-whistle annotations needed to train detectors. RidgeTrace needs none: a small convolutional network (14,209 parameters), trained only on synthetic whistles, clicks and sweeps added to real ocean noise, estimates the probability that each spectrogram cell lies on a whistle, and a rule-based tracker links these cells into contours. Without retraining, we compared a 5–24 kHz version with Silbido, an established rule-based detector, on 12 DCLDE 2011 recordings of four delphinid species (1,516 scored whistles within 5–24 kHz). Averaged over species, RidgeTrace had higher recall (86.9% vs 72.9%; difference 95% range +4.3 to +22.1) and lower precision (56.3% vs 76.4%; *−*32.2 to *−*2.6); F1 did not differ clearly (67.1 vs 74.6). In a blind sample from the four recordings where either method performed weakest, 45% of RidgeTrace’s sampled false alarms were unannotated whistles or harmonics, fragments or echoes of annotated whistles. On 145.5 min of beaked and sperm whale recordings from the training-noise survey, it produced 0.90 false alarms per minute against Silbido’s 4.90 with both held to Silbido’s 150 ms minimum length (4.46 vs 5.06 before this common filter). Its precision must improve before unreviewed use.

## 1. Introduction

Many toothed whales (odontocetes) produce whistles: narrowband, frequency-modulated tonal sounds that usually last from a few tenths of a second to about a second and fall mostly between a few kilohertz and a few tens of kilohertz. Whistles carry information about the caller. Bottlenose dolphins (*Tursiops truncatus*) develop individually distinctive signature whistles [1, 2] and copy one another’s whistles [3], and whistle contour features differ enough between delphinid species to support acoustic species identification [4–6]. For these uses, a whistle is usually represented by its *contour*, the sequence of its fundamental frequency over time, traced on a spectrogram. Passive acoustic monitoring produces far more recordings than analysts can annotate by hand [7, 8]. Automatic contour extraction has therefore been studied for a long time. Most established methods are rule based: they find spectral peaks that stand out from the background noise and link them across time. Silbido [9] finds energy peaks above a signal-to-noise threshold and links them with a graph search, and it is widely used as a reference. Other approaches include peak tracking in PAMGuard [10, 11], a contour detector for whistles and moans [12], image-processing methods [13, 14], a representative-frequency detector [15], and multi-target tracking with probability hypothesis density filters [16, 17]. These methods need no training, but their simple local tests can confuse whistles with tonal noise and with the broadband clicks and burst pulses that dolphins produce [18], and faint or crossing whistles are hard to follow.

Deep learning has improved the detection of many animal sounds [19], including bird song [20], baleen whale calls [21–23] and sperm whale clicks [24]. For whistles, convolutional networks have been used both to classify whistle spectrograms [25] and to extract contours [26, 27]. The main obstacle is training data: a contour extractor needs examples in which every whistle is traced, and such annotations are expensive and rare. Li et al. [26] addressed this by training on synthetic whistles combined with real backgrounds, and Li et al. [27] trained a contour-tracking network without human-annotated training data and evaluated it on the DCLDE 2011 benchmark [28]. Training on simulated data and testing on real data is a common strategy in other fields too [29, 30].

The detector described here was built for a practical need. We were reproducing the VAE-GMM generative model of cetacean whistles of Park et al. [31], which learns from whistle contours extracted from recordings; the whistle detector and tracker used in that study were not available to us, and our recordings, from a 2017 towed-array survey around the Hawaiian Islands [32], have species and encounter times logged but no whistle-level annotations. We therefore built a replacement that could be trained without any human labels. The system goes beyond detecting that a whistle is present: it localises each whistle in the time–frequency plane and returns its contour. It does not estimate the position of the animal. For brevity, we call the full system a detector. It follows the same general idea as Li et al. [26]: synthetic whistles in real noise. To our knowledge, it differs from that work in several respects that may be useful to others:

(1) the network is very small (14,209 parameters) and keeps the full spectrogram resolution, so it trains in under two hours on a free cloud GPU and runs faster than real time on an ordinary CPU;
(2) the real noise comes from a different survey, region and recording system than the test data, so the benchmark tests generalisation, not memorisation of a recording setup;
(3) probable real whistles hidden in the training noise are found with a simple unsu-pervised rule and removed from the loss, instead of being taught to the network as “not a whistle”;
(4) besides the standard benchmark, we measured false alarms on recordings of species that click but are not known to whistle, compared these with Silbido at three detection thresholds, and inspected samples of each method’s false alarms blind to which method produced them.

We report the results in full, including where RidgeTrace does worse than Silbido and the settings that were chosen after test results were seen (Section 4.4). We do not claim a new state of the art: to our knowledge, the best published model on this benchmark [27] was scored over a wider frequency band and cannot be compared directly with RidgeTrace’s numbers.

## 2. Materials and methods

### 2.1. Overview

The detector has two stages (Figure 1). A neural network, which we call RidgeNet, takes a spectrogram and outputs, for each time–frequency cell, a probability that the cell lies on a whistle. A tracker, which uses fixed rules and no learning, then joins high-probability cells into whistle contours and rejects contours that are too short, too weak, duplicated or otherwise implausible. The same method was trained in two frequency bands: 2–24 kHz, used to build the whistle dataset for our reproduction of the VAE-GMM model [31], and 5–24 kHz, matching the lower limit of the DCLDE 2011 annotations. Only the 5–24 kHz version is evaluated on annotated data in this paper; the 2–24 kHz version was evaluated on synthetic test data and on real whistle-free recordings (Section 2.7), but not on annotated real recordings; Section 3.5 illustrates its output on one 2017 false killer whale recording. Table 1 lists the main settings.

**Figure 1.**
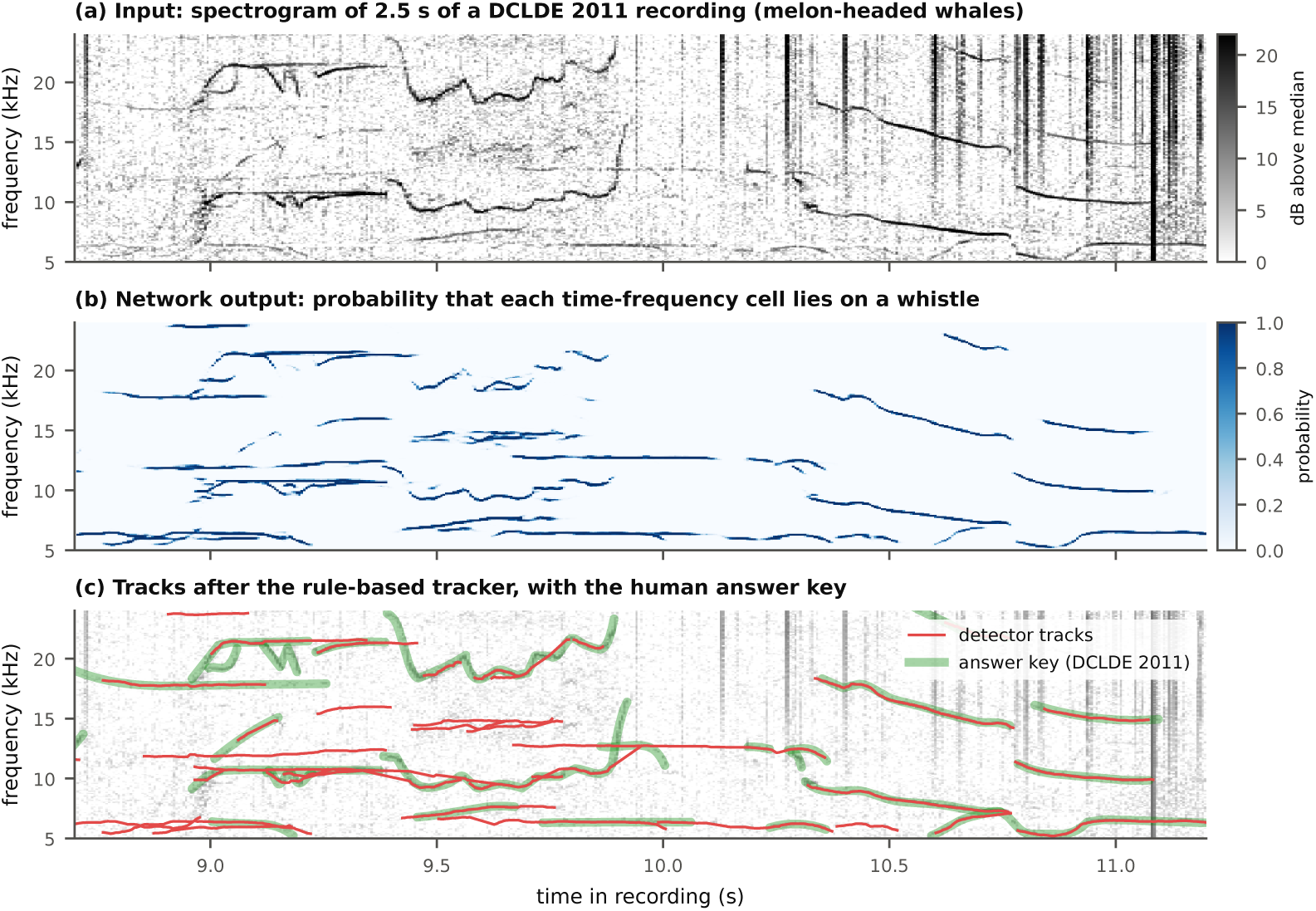
What the detector does, on 2.5 s of a DCLDE 2011 recording of melon-headed whales (`palmyra102006-061020-204327_4`). (a) Input spectrogram, shown as dB above the median of each frequency bin; whistles are thin curved lines, clicks are vertical streaks. (b) Network output: the probability that each time–frequency cell lies on a whistle. Clicks are almost entirely suppressed. (c) Contours after the tracker (red) over the human annotations (green; “answer key” in the legend). The tracks shown are the benchmark detections scored in Section 3.2. Some tracks have no annotation (for example near 12.7 kHz at 9.7–10.2 s); Section 3.4 examines such cases.

**Table 1.** Main settings of the 5–24 kHz detector. All were fixed before the DCLDE 2011 test except the two marked *^∗^*, which apply only to benchmark scoring (Section 4.4).

| Part | Setting | Value |
|---|---|---|
| Spectrogram | Sample rate; high-pass | 192 kHz; 2 kHz, 4th-order Butterworth, zero phase |
|  | Window; hop; resolution | 2,048 samples Hamming; 1,024 samples; 93.75 Hz × 5.33 ms |
|  | Band | 5,062.5–24,000 Hz (203 bins) |
| Conditioning | Floor estimate | Median of 64-frame blocks, running median over 7 blocks |
| Network | Layout | Stem + 6 dilated residual blocks, 16 channels, no pooling |
|  | Parameters; receptive field | 14,209; 39 frames × 19 bins (208 ms × 1.8 kHz) |
| Training | Steps; batch; crop | 6,000; 16; 256 frames (1.37 s) |
|  | Optimiser | AdamW, peak learning rate 0.003, one-cycle schedule, weight decay 10<sup>−4</sup> |
|  | Loss | Weighted binary cross-entropy (positive weight 6) + Dice |
|  | Checkpoint | Best synthetic validation cell-level F1 (step 2,000) |
| Tracker | Peak threshold *τ* | 0.4 (chosen on synthetic validation data) |
|  | Gate; maximum gap | 3 bins + 0.5 per missed frame, at most 8; 12 frames (64 ms) |
|  | Track rejection | <12 frames; <50% observed; mean probability <0.55;<br>mean slope >3 bins/frame; duplicates; synchronous pulse-train combs |
|  | Harmonic removal<sup>*</sup> | Off for the benchmark (on for the 2–24 kHz dataset) |
| Scoring | Minimum detection length<sup>*</sup> | 150 ms (Silbido's default) |

### 2.2. Signal processing

Recordings were read as single channels (channel 0 for multichannel files) and scaled to [*−*1, 1]. They were resampled to 192 kHz with a polyphase filter (Kaiser window, *β* = 5), high-pass filtered at 2 kHz with a fourth-order Butterworth filter applied forwards and backwards (zero phase), and converted to a spectrogram with a 2,048-sample periodic Hamming window [33] (10.67 ms), a hop of 1,024 samples (5.33 ms, 50% overlap) and a 2,048-point FFT. This gives cells of 93.75 Hz by 5.33 ms. Magnitudes were converted to decibels as 20 log_10_(*|X|* + 10*^−^*^12^). The 5–24 kHz detector uses bins 54–256 (5,062.5–24,000 Hz; 203 bins). The 192 kHz rate, 2 kHz high-pass filter and spectrogram settings follow Park et al. [31]; signal processing used SciPy [34].

#### Conditioning

Recording levels differ greatly between recordings and are not calibrated, so the network sees each cell’s level relative to the local noise floor. For each frequency bin, the floor was estimated as the median within blocks of 64 frames (0.34 s), smoothed with a running median over seven blocks (about 2.4 s) and interpolated back to every frame. The network input is

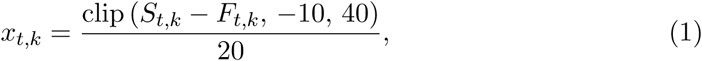

where *S_t,k_*is the spectrogram level in dB at frame *t* and bin *k*, and *F_t,k_* is the floor. Medians are used because whistles are sparse and loud and would bias a mean upwards.

### 2.3. Network

RidgeNet (Figure 2) is a fully convolutional network with no pooling, so its output has exactly the same size as its input and stays aligned cell for cell with the spectrogram. A 3 *×* 3 convolutional stem with 16 channels is followed by six residual blocks [35], each a 3*×*3 dilated convolution [36] with 16 channels, batch normalisation [37] and a ReLU, added to the block’s input. The dilations (time, frequency) of the six blocks are (1,1), (2,1), (4,2), (8,2), (1,1) and (2,1): wider in time than in frequency, because whistles are long and thin. A 1 *×* 1 convolution and a sigmoid give the output probability. The network has 14,209 parameters and its receptive field is 39 frames by 19 bins (208 ms by 1.8 kHz). Long recordings are processed in tiles of 2,048 frames with 32 frames of overlap on each side, which exceeds the receptive-field radius, so tiling does not change the output. The network was implemented in PyTorch [38]. Unlike encoder– decoder segmentation networks such as U-Net [39], which reduce and then restore the resolution, this design never downsamples, which keeps it small and keeps fine frequency detail.

**Figure 2.**
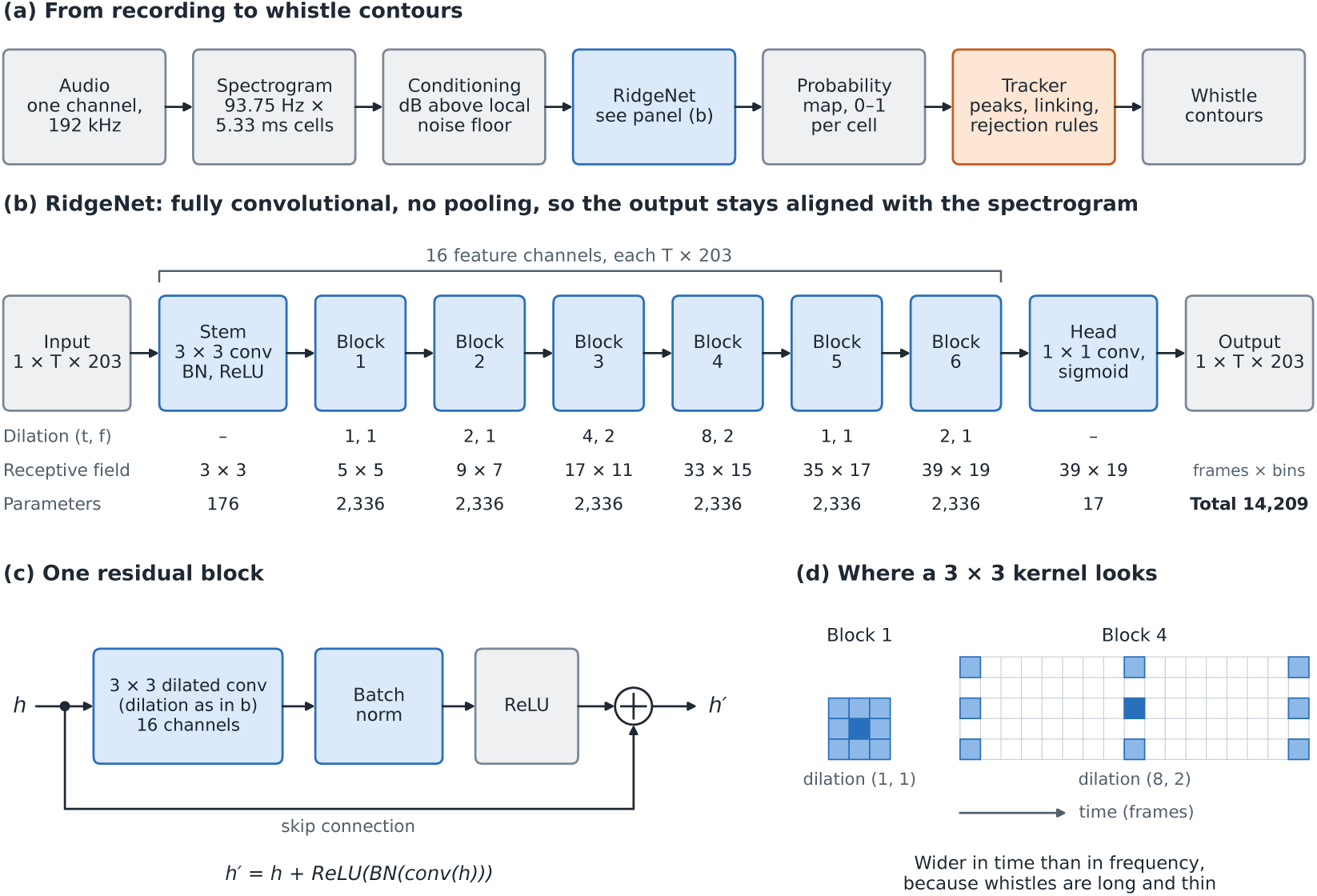
Architecture of the detector. (a) Processing chain, from a recording to whistle contours. Blue marks learned layers, orange the fixed-rule tracker, and grey data and fixed operations. (b) RidgeNet. *T* is the number of time frames and 203 the number of frequency bins in the 5–24 kHz band. The receptive field is the region of the input, in frames *×* bins, that can influence one output cell after each layer. (c) Each residual block adds the output of a dilated convolution, batch normalisation and a ReLU to the block’s input. (d) Cells read by a 3 *×* 3 kernel (light blue) around the current cell (dark blue) in blocks 1 and 4; the vertical axis is frequency, in bins.

### 2.4. Training data

#### Real backgrounds

The noise came from the DCLDE 2022 dataset: recordings from the 2017 Hawaiian Islands Cetacean and Ecosystem Assessment Survey (HICEAS), made aboard the NOAA ship *Reuben Lasker* with a six-channel towed hydrophone array (HTI-96-min hydrophones) sampled at 500 kHz and stored as 60-s files [32, 40]. We used 56 files from a single false killer whale (*Pseudorca crassidens*) encounter (encounter 191), divided into 24 training, 16 validation and 16 test files with no file in more than one split. Each file was cut into 3-s candidate windows starting every second, and windows were ranked by a simple ridge score: the fraction of frames containing a narrow (neighbours 3–5 bins away not raised), persistent (*≥*70% of a 9-frame span) peak at least 8 dB above the floor. The least whistle-like windows with a score of at most 0.02 were kept, up to 20 (training) or 12 (validation and test) per file, with at most 50% overlap. Because the encounter was rich in whistles, clean windows were scarce: 180 training, 44 validation and 116 test windows passed. Half of the training windows were played backwards.

#### Synthetic whistles

Each window received 0–5 synthetic whistles (probabilities 0.10, 0.25, 0.25, 0.18, 0.12 and 0.10; mean 2.27). A whistle’s duration was drawn log-uniformly from 0.1–1.0 s, and its contour was built from 2–6 control points according to one of six shapes (rising 25%, wavy 25%, falling 20%, arch 15%, valley 10%, flat 5%) and interpolated with a shape-preserving piecewise cubic [41], with an optional sinusoidal ripple (probability 0.3; 5–20 Hz rate, 30–250 Hz depth). Contours were kept within 5.3–23.7 kHz and below 60 kHz/s. Second whistles were placed at random (60%), crossing the first (20%) or parallel to it 2–4 bins away (20%), so that the network learned to keep crossing and neighbouring whistles separate. Loudness was set relative to the local background level at the whistle’s frequency, with signal-to-noise ratios (SNR) drawn from *−*2–4 dB (25%), 4–12 dB (50%) and 12–24 dB (25%), so that most whistles were faint. Second and third harmonics were added with probabilities 0.6 and 0.3, at 3–15 dB below the fundamental. Waveforms were synthesised from the running phase of the contour, with tapered onsets and offsets.

#### Distractors

To teach the network to ignore impulsive sounds, 60% of windows received 1–3 synthetic click trains (3–25 clicks, 10–80 ms apart, centre frequencies 8– 70 kHz, decay 0.05–0.30 ms, 10–35 dB above the window’s RMS level), and 20% received trains of broadband sweeps resembling echosounder pings. Distractors were never labelled as targets.

#### Targets and weights

The target for each cell was a soft ridge centred on the true frequency of each synthetic whistle, 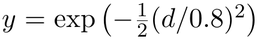, where *d* is the distance in bins; harmonics were targets as well. Each cell also had a weight *w*. Cells on parts of a whistle with SNR below 3 dB, and cells within 2 bins of them, received *w* = 0: the network was neither rewarded nor penalised there, because those parts are not visible. All other cells had *w* = 1.

#### Ignoring real whistles in the background

Screening cannot remove every real whistle from the noise, and an unscreened faint whistle would be taught as “not a whistle”. We therefore marked probable real ridges in every background window with a classical rule: for each cell, the median level above the floor was computed along 25-frame line segments through the cell at 13 slopes between *−*1 and +1 bin per frame, and the cell was marked if the largest of these medians exceeded 4 dB. Marked cells (dilated by 1 frame and 3 bins) received *w* = 0. A median along a line is high only if most of the line is raised, so clicks and random noise are rarely marked. The rule marked 2.19% of cells in the real backgrounds and 0.65% in phase-randomised copies of them, which suggests that most of what it marks is real tonal structure. In a development comparison on synthetic data, adding this rule to the 2–24 kHz detector’s training raised whistle-level recall on the 617 whistles of the synthetic validation set from 73.9% to 76.0%. Of the 35 whistles found by only one of the two versions, 24 were found only with the rule and 11 only without it (exact McNemar test, *p* = 0.041); the gain was largest among faint whistles (below 5 dB, recall 5.6% to 12.7%).

#### Loss and optimisation

The loss was a weighted binary cross-entropy plus a soft Dice term [42]:

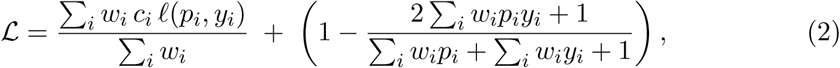

where *p_i_*is the network output at cell *i*, *ℓ* is the binary cross-entropy, and *c_i_* = 6 for whistle cells (*y_i_ >* 0.1) and 1 otherwise, to offset the rarity of whistle cells. Training ran for 6,000 steps with batches of 16 freshly generated 256-frame crops (96,000 examples in total), using AdamW [43] with weight decay 10*^−^*^4^ and a one-cycle learning-rate schedule [44] peaking at 0.003. Every 500 steps the network was scored on a fixed synthetic validation set of 600 examples, and the checkpoint with the best cell-level F1 was kept; this was step 2,000 (F1 0.559; F1 varied only between 0.533 and 0.559 after step 500). Training took 101 min on a free Google Colab NVIDIA Tesla T4 GPU (Python 3.13, PyTorch 2.11). The 2–24 kHz detector was trained in the same way, with synthetic whistles between 2.3 and 23.7 kHz and its own background windows selected in its band; its best checkpoint was at step 1,500, training took about 65 min on an NVIDIA RTX 5060 laptop GPU, and its peak threshold is 0.5.

### 2.5. Tracker

#### Peaks

In each frame, local maxima of the network output at or above the threshold *τ* were taken as peaks and refined to a fraction of a bin as *k^′^* = *k* +(*p_k_*_+1_ *− p_k−_*_1_)*/*(*p_k−_*_1_ + *p_k_* + *p_k_*_+1_). Peaks less than 2 bins apart were merged, keeping the stronger. Frames in which more than 30% of the band exceeded *τ* were treated as broadband (clicks) and skipped.

#### Linking

Active tracks were extended frame by frame. Each track’s next frequency was predicted by a straight line through its last five points, and a peak could join the track only if it fell within a gate of min(8, 3 + 0.5*m*) bins of the prediction, where *m* is the number of consecutive frames the track had gone without a peak. The cost of assigning peak *j* to track *i* was 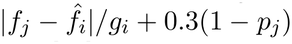, and peaks were assigned to tracks by the Hungarian algorithm [45], which minimises the total cost. Unassigned peaks started new tracks; tracks without a peak for more than 12 frames (64 ms) were ended. This is standard gated nearest-neighbour multi-target tracking [46]; the linear prediction lets tracks pass through crossings and short gaps, which were filled by interpolation and flagged as not observed.

#### Rejection

A track was rejected if it spanned fewer than 12 frames (64 ms), if fewer than half of its frames had an observed peak, if its mean network output was below 0.55, or if its mean slope exceeded 3 bins per frame (52.7 kHz/s). A track that followed the path of a stronger track within 1.5 bins for at least 80% of its length was rejected as a duplicate, and six or more tracks starting within 3 frames of each other with similar lengths were rejected as a pulse-train artefact. Optionally, tracks at 2, 3 or 4 times the frequency of a lower, overlapping track (within 3%) were rejected as harmonics. Harmonic removal was used to build the VAE-GMM reproduction dataset, which needs fundamental contours only, but not for the benchmark, because the DCLDE 2011 annotations include harmonics.

#### Threshold

The peak threshold was chosen on synthetic validation data only. Over *τ* = 0.30, 0.35*, . . . ,* 0.80, we kept the values that produced at most 2 false tracks per minute on “clean negatives” (phase-randomised backgrounds, which keep the noise spectrum but cannot contain whistles) and whose whistle-level recall was within 0.01 of the best, and chose the one nearest 0.5. This gave *τ* = 0.4. Validation recall varied by only about 0.04 across the whole grid, so the exact value matters little.

### 2.6. Evaluation on DCLDE 2011

#### Recordings

We used the test set of Li et al. [27] from the DCLDE 2011 workshop data [28]: 12 single-channel recordings sampled at 192 kHz, three each of bottlenose dolphin, long-beaked common dolphin (*Delphinus capensis* in the dataset documentation; now usually treated as *D. delphis bairdii* ), melon-headed whale (*Peponocephala electra*) and spinner dolphin (*Stenella longirostris*), totalling 43.2 min and 6,011 annotated whistles (Appendix A). The detector was not trained or tuned on DCLDE 2011 data; two post-processing choices were made after the first test run (Section 4.4). The species, regions and recording equipment all differ from those of the training noise. All files were checked against their published sizes and checksums.

#### Scoring

We re-implemented the scoring procedure in Python from Silbido’s scoring code, following the rules of Li et al. [27]. An annotated whistle was counted (expected to be found) if it lasted at least 150 ms, was more than 10 dB above the noise over at least a third of its length and, in our main scoring, lay entirely within 5–24 kHz; 1,516 of the 6,011 annotated whistles met these criteria. We used the original 2011 annotations throughout, and also scored RidgeTrace against the revised annotations released with the dataset in 2025, which extend the annotations to weaker whistles in a small set of files [28]. A detection matched an annotated whistle if their mean frequency difference over their time overlap was at most 350 Hz. Detections that matched only uncounted whistles were ignored, counting neither as correct nor as false. In 5–24 kHz scoring, detections whose median frequency lay outside the band were also ignored, so that Silbido, which searches a wider band, was not penalised. Precision is the share of the remaining detections that matched a counted whistle, and recall is the share of counted whistles matched by at least one detection. Counts were pooled within each species, and precision, recall and F1 were then averaged across the four species. We also report the mean frequency deviation of matched detections, their coverage (the share of the whistle’s duration they span) and fragmentation (matched detections per found whistle). The scorer was checked by scoring the annotations against themselves (recall 100%, precision 99.93%) and against shifted and shortened copies, which were rejected as expected.

#### The 30% rule

Li et al. [27] state that a detection must also cover more than 30% of the annotated whistle, but we found no such rule in Silbido’s scoring code, and our Silbido scores agree much more closely with their published Silbido scores without it (Section 3.2). We therefore report both versions and treat the version without the 30% rule as the main score. We also report scores counting whistles at all frequencies, since the 5–24 kHz criterion excludes 344 of the 1,860 whistles that would otherwise count (18%).

#### Silbido

Silbido 3.0 [9] was run in MATLAB R2024b with its classic graph search and default settings (energy peaks at least 10 dB above the noise; minimum whistle length 150 ms) on the same 12 recordings. The run was later repeated on a different computer and operating system: 10 of the 12 output files were identical, two differed slightly, and all scores were identical.

#### Benchmark settings for RidgeTrace

For the benchmark, harmonic removal was switched off and detections shorter than 150 ms (Silbido’s default minimum) were discarded. Both choices were made after seeing test results (Section 4.4); we report the effect of each.

#### Statistics

Uncertainty was estimated with a bootstrap [47]: 10,000 resamples, each drawing three recordings per species with replacement and recomputing the species-averaged score. The same resamples were used for both methods, so the ranges of the differences are paired. With only three recordings per species these ranges are approximate. Differences between methods were also tested with exact two-sided Wilcoxon signed-rank tests [48] on the 12 per-recording values (ties dropped). Because three metrics (precision, recall and F1) were tested on the same recordings, *p* values were adjusted for multiple comparisons with the Holm method [49] across the three metrics within each scoring version. We report adjusted values in the text and unadjusted values in the tables. Differences were computed from unrounded values, so they can differ by 0.1 from the difference of the rounded numbers shown. The 12 recordings fall into four species groups of three, so they are not fully independent, and all *p* values should be read as approximate.

### 2.7. Whistle-free recordings

The DCLDE 2011 analysts were asked to annotate strong whistles, and many very short whistles were skipped [28], so a false alarm on that benchmark may be a real whistle. To measure false alarms when no whistles are expected, we ran both methods on recordings of species that produce clicks but are not generally reported to whistle: beaked whales, which echolocate with frequency-modulated clicks [50, 51], and sperm whales (*Physeter macrocephalus*), which produce clicks and codas [52]. (Whistle-like sounds have occasionally been recorded near beaked whales [53, 54]; we return to this in Section 4.)

Before running any detector, we selected from the HICEAS 2017 encounter log [32] the 21 single-species encounters with no whistling species logged within 15 min: 13 beaked whale encounters (Cuvier’s, *Ziphius cavirostris*, and Blainville’s, *Mesoplodon densirostris*) and 8 sperm whale encounters. We used every 1-min file overlapping a beaked whale encounter (72 files, 71.5 min) and up to 10 files spread evenly through each sperm whale encounter (74 files, 74.0 min): 146 files and 145.5 min in total, listed with checksums before analysis. These recordings come from the same survey and equipment as RidgeTrace’s training noise, though from different encounters, so this test is not fully independent of training.

Every detection counted as a false alarm. Both RidgeTrace versions ran with their fixed settings: the 5–24 kHz detector as on the benchmark, and the 2–24 kHz detector with the settings used to build the dataset (peak threshold 0.5, harmonic removal on). Silbido ran at its default settings on 16-bit WAV copies of the same audio (channel 0, resampled to 192 kHz; 1,073 of about 1.7 billion samples were clipped, from the loudest clicks). We counted detections under the same rule as the benchmark (median frequency within 5–24 kHz, at least 150 ms long) and, separately, all detections each method produced. Ranges are 95% bootstrap ranges over encounters (10,000 resamples), paired between methods.

#### Silbido’s threshold

To see whether a different setting of Silbido would match RidgeTrace, Silbido was also run at peak thresholds of 8 and 12 dB, values fixed before the run, on both the DCLDE 2011 recordings and the whistle-free recordings.

### 2.8. What the false alarms are

To understand the false alarms, we sampled them and looked at them. On the four recordings where either method was weakest (RidgeTrace’s three lowest-precision recordings and Silbido’s two lowest-F1 recordings, one of which is shared: `palmyra092007FS192-070925-023000, Qx-Tt-SCI0608, Qx-Dc-CC0411` and `palmyra092007FS192-071004-032342`), up to 20 false alarms per method per recording were drawn at random with a fixed seed under the main scoring (147 in total: 80 RidgeTrace’s, 67 Silbido’s). Each was drawn as a spectrogram picture with the annotations overlaid, and the pictures were shuffled and numbered so that the person sorting them could not tell which method produced each one. One person, the corresponding author (B. S. Parjanya), sorted them into six categories: real whistle not in the annotations (W); harmonic, piece or echo of an annotated whistle (H); tonal noise such as an engine or echosounder (T); clicks or burst pulses (C); other noise (N); and can’t tell (?), and marked each decision as sure or unsure. Whether the blinding succeeded was not tested. The same procedure was applied to 10 random detections per detector per species group in the whistle-free test (40 pictures for the two RidgeTrace versions) and to 20 of Silbido’s whistle-free detections, which were mixed with 10 of the earlier pictures shown again, on a later day, to check consistency. These checks describe samples; they do not change any score.

## 3. Results

### 3.1. Synthetic test

On synthetic examples built from the 16 held-out test noise files, the 5–24 kHz detector found 63% of the synthetic whistles (338 of 537; a whistle counted as found if one track covered at least 60% of its visible frames within 1.5 bins). Recall depended strongly on loudness (Table 2): the detector was nearly blind below 5 dB SNR and reliable above 10 dB. Neither detector produced any false track on clean negatives. The 2–24 kHz detector had similar recall at each SNR; its higher overall recall (0.729) reflects a test mix with fewer faint whistles. These numbers describe the synthetic task only, not performance on real recordings.

**Table 2.**
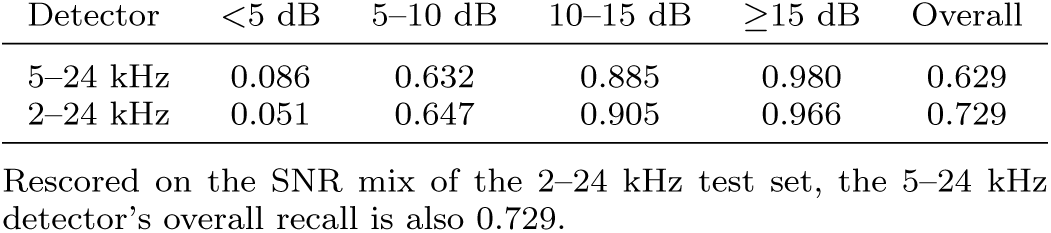
Recall on synthetic test data by whistle signal-to-noise ratio. The two test sets were generated separately, so the comparison is not paired.

| Detector | <5 dB | 5–10 dB | 10–15 dB | ≥15 dB | Overall |
|---|---|---|---|---|---|
| 5–24 kHz | 0.086 | 0.632 | 0.885 | 0.980 | 0.629 |
| 2–24 kHz | 0.051 | 0.647 | 0.905 | 0.966 | 0.729 |

Rescored on the SNR mix of the 2–24 kHz test set, the 5–24 kHz detector’s overall recall is also 0.729.

### 3.2. DCLDE 2011

Averaged over species, RidgeTrace found 86.9% of the counted whistles and Silbido 72.9%, a difference of +14.0 percentage points (95% range +4.3 to +22.1; Holm-adjusted *p* = 0.041; Table 3). Silbido was more precise: 76.4% of its detections were correct against 56.3% of RidgeTrace’s (*−*20.2, range *−*32.2 to *−*2.6; adjusted *p* = 0.085).

**Table 3.** DCLDE 2011 test set (12 recordings, 4 species), 5–24 kHz scoring, detections *≥*150 ms. Species average with 95% bootstrap range; the difference is paired. *p*: Wilcoxon signed-rank test on the 12 recordings, unadjusted (Holm-adjusted across the three metrics, no 30% rule: precision 0.085, recall 0.041, F1 0.176; with 30% rule: 0.154, 0.037, 0.380).

| Scoring | Metric | RidgeTrace | Silbido | RidgeTrace − Silbido | *p* |
|---|---|---|---|---|---|
| No 30% rule (main) | Precision | 56.3 [46.8, 72.2] | 76.4 [62.2, 91.2] | −20.2 [−32.2, −2.6] | 0.042 |
|  | Recall | 86.9 [83.6, 91.0] | 72.9 [64.8, 83.0] | +14.0 [+4.3, +22.1] | 0.014 |
|  | F1 | 67.1 [58.3, 78.8] | 74.6 [63.0, 86.4] | −7.5 [−18.0, +5.9] | 0.176 |
| With 30% rule | Precision | 47.1 [39.2, 61.1] | 64.5 [52.4, 76.8] | −17.4 [−27.9, −1.1] | 0.077 |
|  | Recall | 85.2 [81.0, 89.8] | 70.0 [62.0, 80.2] | +15.1 [+4.8, +23.5] | 0.012 |
|  | F1 | 59.9 [51.7, 71.1] | 67.1 [56.7, 77.8] | −7.2 [−17.3, +6.1] | 0.380 |

The bootstrap range of the precision difference excluded zero, but after adjustment for three tests the Wilcoxon test alone did not reach *p <* 0.05, so we treat both differences as clear in direction but approximate in size. Silbido’s F1 was higher by 7.5 points (74.6 vs 67.1), but the range of the difference included zero (*−*18.0 to +5.9; adjusted *p* = 0.18), so on this test set neither method was clearly better overall. The pattern was the same with the 30% rule: all scores were lower, by about 9 points for RidgeTrace’s precision and 12 for Silbido’s, while the gaps between methods hardly changed. Matched detections of the two methods had similar frequency accuracy (mean deviation 94 vs 100 Hz, well below the 93.75 Hz cell size), and RidgeTrace’s covered somewhat more of each whistle (86.6% vs 80.7%; no uncertainty estimated), with slightly more fragmentation (1.30 vs 1.25 detections per found whistle).

Results varied between species and recordings (Table 4). RidgeTrace had the higher recall on 9 recordings (1 tie) and Silbido the higher precision on 10. Sil-bido had the higher F1 on 9 recordings and RidgeTrace on 3 (one of Silbido’s, `palmyra092007FS192-071011-232000`, by 84.18 to 84.17). By species, Silbido had the higher F1 for bottlenose dolphins, melon-headed whales and spinner dolphins, and RidgeTrace for common dolphins, largely because of one recording (`Qx-Dc-CC0411`) on which Silbido found only 21% of the whistles. RidgeTrace’s lowest precision was on `palmyra092007FS192-070925-023000` (18.0%), a very busy melon-headed whale recording in which only 76 of 1,067 annotated whistles met the scoring criteria; Section 3.4 examines its false alarms.

**Table 4.** Results by species and recording (5–24 kHz, no 30% rule). Counted: annotated whistles meeting the scoring criteria, out of all annotated whistles. Full recording names are in Appendix A. Higher F1 in bold.

| Species | Recording | Counted / annotated | Silbido P / R / F1 | RidgeTrace P / R / F1 |
|---|---|---|---|---|
| Bottlenose dolphin | SCI0608 | 56 / 152 | 74.4 / 85.7 / **79.7** | 35.0 / 85.7 / 49.7 |
|  | 070924-205305 | 99 / 539 | 97.1 / 83.8 / **90.0** | 86.4 / 81.8 / 84.1 |
|  | 070924-205730 | 103 / 336 | 93.9 / 80.6 / 86.7 | 89.1 / 91.3 / **90.2** |
|  | *Species* | 258 / 1,027 | 89.6 / 82.9 / **86.1** | 65.3 / 86.4 / 74.4 |
| Common dolphin | CC0411 | 287 / 759 | 26.4 / 21.3 / 23.5 | 35.4 / 81.2 / **49.3** |
|  | FLIP0610 | 219 / 533 | 93.2 / 89.5 / **91.3** | 65.3 / 94.1 / 77.1 |
|  | SC03 | 9 / 135 | 90.9 / 77.8 / **83.8** | 57.1 / 100 / 72.7 |
|  | *Species* | 515 / 1,427 | 58.4 / 51.3 / 54.6 | 45.0 / 87.0 / **59.3** |
| Melon-headed whale | 071004-032342 | 54 / 202 | 34.1 / 70.4 / 46.0 | 62.5 / 94.4 / **75.2** |
|  | 061020-204327_4 | 153 / 601 | 94.0 / 75.2 / **83.5** | 71.2 / 83.7 / 76.9 |
|  | 070925-023000 | 76 / 1,067 | 89.4 / 69.7 / **78.4** | 18.0 / 90.8 / 30.1 |
|  | *Species* | 283 / 1,870 | 71.3 / 72.8 / **72.0** | 38.5 / 87.6 / 53.5 |
| Spinner dolphin | 070927-224737 | 237 / 734 | 91.6 / 86.5 / **89.0** | 77.0 / 92.4 / 84.0 |
|  | 061103-213127_4 | 112 / 553 | 76.1 / 85.7 / **80.6** | 68.2 / 77.7 / 72.6 |
|  | 071011-232000 | 111 / 400 | 88.6 / 80.2 / **84.2** | 84.6 / 83.8 / 84.2 |
|  | *Species* | 460 / 1,687 | 86.4 / 84.8 / **85.6** | 76.2 / 86.7 / 81.1 |
| Species average |  | 1,516 / 6,011 | 76.4 / 72.9 / **74.6** | 56.3 / 86.9 / 67.1 |

#### Other scoring choices and settings

When whistles at all frequencies were counted, RidgeTrace’s recall fell to 76.6% (Table 5), because RidgeTrace cannot see whistles above 24 kHz, and the recall advantage over Silbido was no longer clear (+4.1, range *−*6.6 to +13.3; unadjusted and Holm-adjusted *p* = 0.97; in that scoring version the unadjusted *p* values for precision, recall and F1 were 0.092, 0.970 and 0.176, and the adjusted values 0.277, 0.970 and 0.353). Without the 150 ms minimum detection length, RidgeTrace found more whistles but made many more false alarms: with the 30% rule, recall rose from 85.2% to 93.3%, but precision fell from 47.1% to 25.6% and F1 from 59.9 to 39.3. Keeping harmonics raised recall by 6.0 points and precision by 0.6 points (with the 30% rule). The revised 2025 version of the annotations changed RidgeTrace’s scores by less than one point.

**Table 5.** Sensitivity to scoring choices and settings. Species-averaged precision / recall / F1 (%). The three rows for RidgeTrace alone are scored with the 30% rule, so compare them with the “5–24 kHz, with 30% rule” row (47.1 / 85.2 / 59.9), not with the main row.

| Scoring or setting | Silbido | RidgeTrace |
|---|---|---|
| 5–24 kHz, no 30% rule (main) | 76.4 / 72.9 / 74.6 | 56.3 / 86.9 / 67.1 |
| 5–24 kHz, with 30% rule | 64.5 / 70.0 / 67.1 | 47.1 / 85.2 / 59.9 |
| All frequencies, no 30% rule | 71.4 / 72.4 / 71.6 | 57.6 / 76.6 / 63.8 |
| All frequencies, with 30% rule | 61.3 / 69.8 / 65.1 | 48.5 / 74.5 / 57.3 |
| *RidgeTrace, 5–24 kHz, with 30% rule:* |  |  |
| No 150 ms minimum detection length | – | 25.6 / 93.3 / 39.3 |
| Harmonics removed | – | 46.5 / 79.2 / 57.9 |
| Revised (2025) annotations | – | 47.9 / 85.2 / 60.5 |

#### Our Silbido run against published results

To check our Silbido run and scorer, we compared them with the Silbido results of Li et al. [27], counting whistles at all frequencies and without the 30% rule (Table 6). Recall, frequency deviation, coverage and fragmentation agreed closely, but our precision was about 10 points lower, mainly for common dolphins (53.0% vs 75.4%) and melon-headed whales (55.9% vs about 66.7%, calculated from their averages). With the 30% rule our fragmentation would be 1.10 rather than their 1.23, which is why we believe the published Silbido scores were computed without that rule. We could not explain the precision gap. It is concentrated in two species rather than spread evenly, so it is unlikely to be a uniform offset of our scorer; it may instead come from differences in particular recordings, in Silbido’s version or settings, or in how individual detections were matched. Both methods were scored by the same scorer under the same rules, but because the cause is unknown we cannot assume it affects them equally, and the absolute precision values in this paper, for both methods, should be read with this in mind. Running our scorer on the detection files behind the published results would isolate the cause. For context only, Li et al. [27] reported precision 88.78%, recall 85.78% and F1 87.20 for their best network, scored over 5–50 kHz; those numbers are not directly comparable with RidgeTrace’s.

**Table 6.** Our Silbido run compared with the Silbido results published by Li et al. [27]: all frequencies, no 30% rule.

|  | Precision | Recall | F1 | Deviation (Hz) | Coverage (%) | Fragmentation |
|---|---|---|---|---|---|---|
| Our run | 71.4 | 72.4 | 71.6 | 101 | 80.5 | 1.23 |
| Li et al. [27] | 81.13 | 72.28 | 75.95 | 101 | 81.05 | 1.23 |

### 3.3. Whistle-free recordings

On the 145.5 min of beaked whale and sperm whale recordings, RidgeTrace (5–24 kHz) produced 131 detections under the benchmark rule, 0.90 per minute, against Silbido’s 713, 4.90 per minute (Table 7). The paired difference was *−*4.00 per minute (95% range *−*5.63 to *−*2.50), and it was similar for beaked whales and for sperm whales. The 2–24 kHz detector, with its dataset settings, produced 0.70 per minute. Before this common benchmark filter, the rates of detections returned under each method’s own operating settings were 4.46 per minute for RidgeTrace (5–24 kHz), 7.61 for the 2–24 kHz detector and 5.06 for Silbido. These are not all-length or frequency-matched rates. Silbido’s default settings already discard detections shorter than 150 ms (none of its 736 whistle-free detections was shorter), and 23 of them lay above 24 kHz, outside RidgeTrace’s band; the output of both RidgeTrace versions includes tracks as short as 64 ms. The benchmark filter therefore removed only short tracks from RidgeTrace (5–24 kHz) (518 of 649), and only the 23 out-of-band detections from Silbido (for the 2–24 kHz detector it also removed tracks with a median frequency below 5 kHz). The difference between the two benchmark-rule rates thus comes almost entirely from applying to RidgeTrace the 150 ms minimum that Silbido applies internally. Both methods are held to the same minimum, but this minimum is one of the two settings chosen after the first DCLDE 2011 results (Section 4.4). False alarms were bursty: a few files and encounters held most of them, hence the wide ranges.

**Table 7.** Whistle-free test: false alarms per minute on 2017 recordings of beaked whales (71.5 min, 13 encounters) and sperm whales (74.0 min, 8 encounters), benchmark rule (median frequency 5–24 kHz, *≥*150 ms). 95% bootstrap range over encounters; number of detections in parentheses after the range for the pooled column.

| Method | Beaked whales | Sperm whales | Both |
|---|---|---|---|
| RidgeTrace, 5–24 kHz | 1.02 [0.47, 1.73] | 0.78 [0.34, 1.23] | 0.90 [0.54, 1.30] (131) |
| RidgeTrace, 2–24 kHz (dataset settings) | 0.81 [0.47, 1.22] | 0.59 [0.29, 0.89] | 0.70 [0.47, 0.94] (102) |
| Silbido, default (10 dB) | 5.21 [2.81, 8.18] | 4.59 [2.06, 7.20] | 4.90 [3.14, 6.81] (713) |
| RidgeTrace (5–24 kHz) − Silbido, paired | −4.19 [−6.58, −2.15] | −3.81 [−6.21, −1.65] | −4.00 [−5.63, −2.50] |

For comparison, under the same frequency and length rule RidgeTrace produced 35.6 false alarms per minute on DCLDE 2011 (1,538 in 43.2 min) and Silbido 10.3 (447). RidgeTrace’s false-alarm rate was therefore about 40 times higher when dolphins were whistling than when no whistling species was present, whereas Silbido’s was about twice as high. The DCLDE 2011 counts exclude detections that matched only uncounted whistles, while the whistle-free counts include every detection, so the DCLDE 2011 rates, and hence these ratios, are if anything underestimates.

#### Silbido’s threshold

Lowering Silbido’s threshold to 8 dB raised its recall on DCLDE 2011 to 80.5% but its whistle-free false-alarm rate to 30.35 per minute; raising it to 12 dB brought the rate down to 0.78 per minute, close to RidgeTrace’s, but its recall fell to 57.0% (Table 8; Figure 3). Silbido’s default was its best setting by F1. At none of the three thresholds did Silbido have both RidgeTrace’s recall and its false-alarm rate. Silbido’s precision on DCLDE 2011 stayed higher than RidgeTrace’s at all three thresholds.

**Figure 3.**
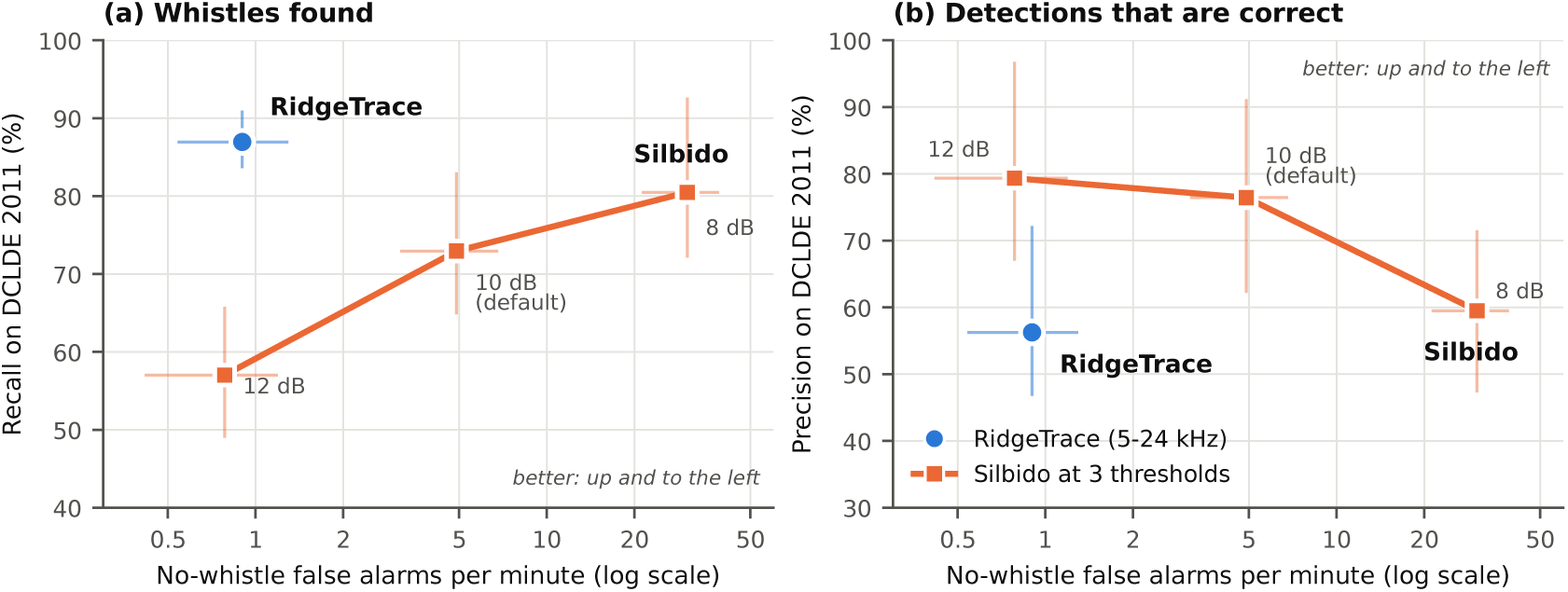
Recall (a) and precision (b) on DCLDE 2011 (5–24 kHz, no 30% rule) against false alarms per minute on the whistle-free recordings (“no-whistle” on the axes; log scale), all under the benchmark rule (median frequency 5–24 kHz, *≥*150 ms), for RidgeTrace (5–24 kHz; blue circle) and Silbido at peak thresholds of 8, 10 (default) and 12 dB (orange squares). Bars show 95% bootstrap ranges: over recordings for DCLDE 2011 and over encounters for the whistle-free test. Better is up and to the left.

**Table 8.**
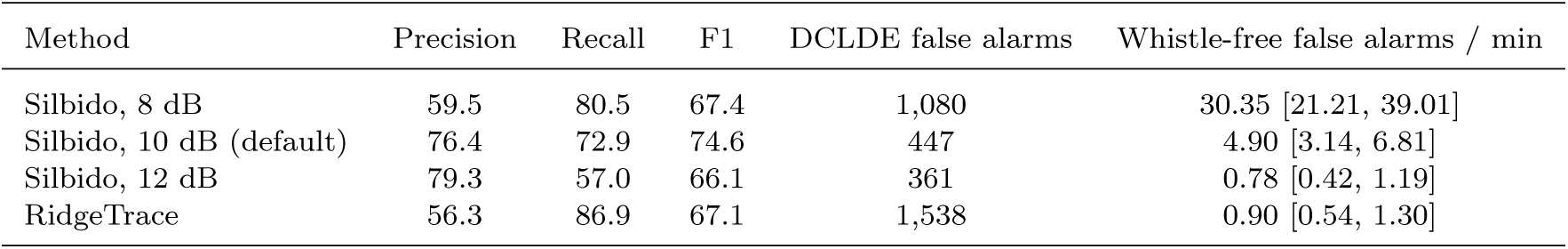
Silbido at three peak thresholds (8, 10 = default, 12 dB; values fixed before the run) and RidgeTrace (5–24 kHz). DCLDE 2011 main scoring; DCLDE false alarms are counts in 43.2 min; whistle-free false alarms per minute with 95% range, benchmark rule.

### 3.4. What the false alarms are

In the blind sample from the four weakest DCLDE 2011 recordings, 24% of RidgeTrace’s sampled false alarms were judged to be real whistles missing from the annotations and 21% harmonics, pieces or echoes of annotated whistles, against 6% and 16% of Silbido’s (Table 9; examples in Figure 4). Silbido’s sampled false alarms were mostly other noise (52%) and clicks or burst pulses (19%). Tonal noise such as echosounders was rare for both methods (2% and 0%). On palmyra092007FS192-070925-023000, where RidgeTrace’s precision was 18%, 15 of the 20 sampled false alarms were unannotated whistles or pieces of annotated ones; on the other three recordings the share was 21 of 60 (35%). Of the 80 decisions on RidgeTrace’s false alarms, 40 were marked sure. Only 84 of the 147 decisions were marked sure, and the sample, at most 20 per method per recording, is not proportional to all false alarms (1,538 RidgeTrace’s and 447 Silbido’s), so these shares describe the sample rather than all false alarms.

**Figure 4.**
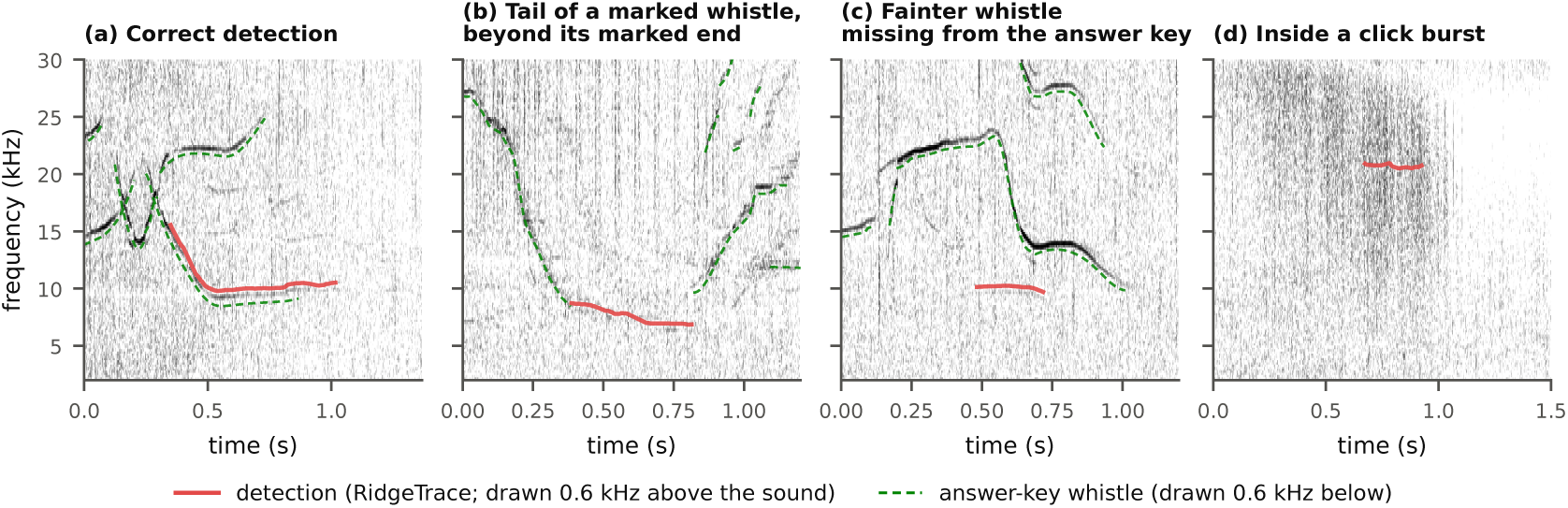
Examples of RidgeTrace’s output on DCLDE 2011 melon-headed whale recordings. Detections (red) are drawn 0.6 kHz above the sound and annotations (green, dashed; “answer-key whistle” in the legend, and “marked” in the panel titles means annotated) 0.6 kHz below it, so that both remain visible. (a) A correct detection, chosen by a fixed rule (071004-032342). (b) A detection along the tail of an annotated whistle, beyond its annotated end, scored as a false alarm (070925-023000). (c) A fainter whistle missing from the annotations, scored as a false alarm (071004-032342). (d) A false alarm inside a burst of clicks (071004-032342). Examples (b)–(d) come from the blind sample and were sorted “sure”.

**Table 9.** Sampled false alarms sorted blind (method hidden) by one person. W: real whistle not in the annotations (whistlefree test: unlogged whistler); H: harmonic, piece or echo of an annotated whistle; T: tonal noise; C: clicks or burst pulses; N: other noise; ?: can’t tell. Samples are not proportional to all false alarms. Percentages are of *n* and rounded to whole numbers (for example, 2 of 80 = 2.5% is shown as 2%), so a row may not sum to exactly 100%.

| Test | Method | *n* (sure) | W | H | T | C | N | ? |
|---|---|---|---|---|---|---|---|---|
| DCLDE 2011 | RidgeTrace | 80 (40) | 19 (24%) | 17 (21%) | 2 (2%) | 8 (10%) | 27 (34%) | 7 (9%) |
|  | Silbido | 67 (44) | 4 (6%) | 11 (16%) | 0 (0%) | 13 (19%) | 35 (52%) | 4 (6%) |
| Whistle-free | RidgeTrace (5–24 kHz) | 20 (12) | 2 (10%) | – | 0 (0%) | 8 (40%) | 10 (50%) | 0 (0%) |
|  | Silbido | 20 (19) | 0 (0%) | – | 0 (0%) | 5 (25%) | 15 (75%) | 0 (0%) |

Consistency: of 10 pictures shown again on a later day, 7 received the same category and all 10 the same whistle-or-not judgement.

In the whistle-free test, 2 of 20 sampled detections from RidgeTrace (5–24 kHz) were judged possible whistles, both in beaked whale recordings and both marked unsure; the rest were noise (10) and clicks (8). Of the 2–24 kHz detector’s 20, 16 were noise, 3 clicks and 1 tonal noise. Of Silbido’s 20, 15 were noise and 5 clicks, all of the latter in sperm whale recordings. Of the 10 pictures shown again on a later day, 7 received the same category; the three changes were all from clicks to other noise, and all 10 received the same whistle-or-not judgement.

### 3.5. The 2–24 kHz detector on a 2017 recording

To illustrate the version used to build the VAE-GMM reproduction dataset, we ran it with its dataset settings (peak threshold 0.5, harmonic removal on) on one minute of a false killer whale encounter that was not used in training (encounter 416, 30 November 2017; Figure 5). The minute was chosen in two steps, both stated here. A minute drawn at random beforehand from the 1,051 one-minute files of the seven false killer whale encounters other than encounter 191 turned out to be the first minute of encounter 416 and contained almost no whistling (highest classical ridge score 0.009; 14 contours, 2 of them at least 150 ms long). After seeing this, we used the middle minute of the same encounter instead (file 34 of 66). The 2.5 s window shown in detail was the one with the highest classical ridge score (Section 2.4), a rule that does not use the network.

**Figure 5.**
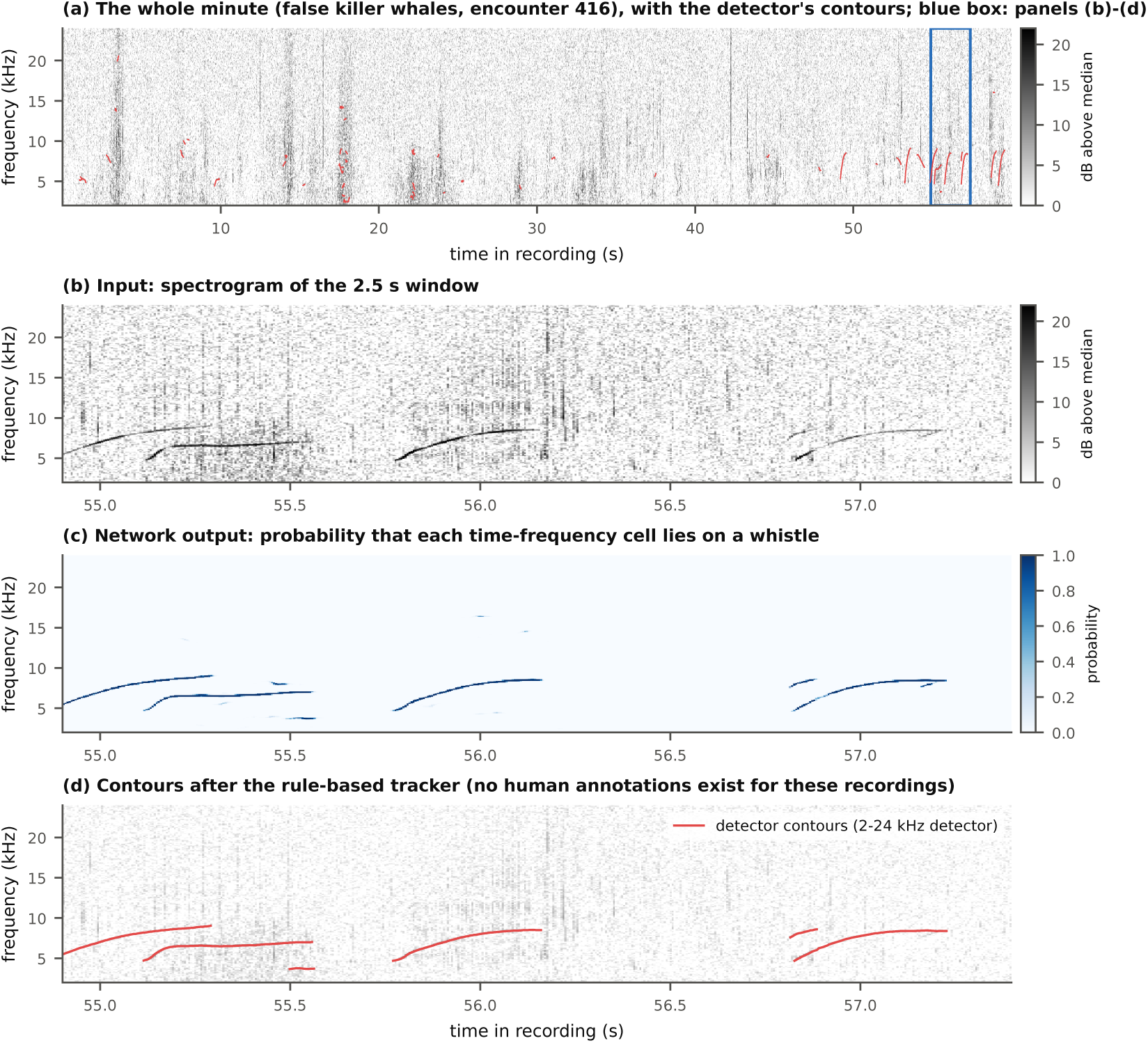
Output of the 2–24 kHz detector, with the settings used to build the VAE-GMM reproduction dataset, on one minute of a 2017 false killer whale recording not used in training (1705 20171130 232045 428, the middle minute of encounter 416; Section 3.5). (a) The whole minute with all 53 kept contours (red); the blue box marks the 2.5 s window shown in (b)–(d), chosen by the highest classical ridge score. (b) Spectrogram, (c) network output and (d) contours in that window. No whistle-level annotations exist for these recordings, so the figure illustrates the detector’s output and does not measure its accuracy.

In that minute the detector kept 53 contours, 17 of them at least 150 ms long; the tracker rejected 69 tracks as too short or mostly interpolated. Judged by eye, four of the six contours in the detailed window follow clearly visible upswept whistles from end to end. One short contour near 3.7 kHz has no visible whistle under it, and one short piece near 8 kHz may belong to a fainter overlapping whistle. Elsewhere in the minute, short contours occur inside click bursts (for example near 17.5 and 22 s), which are probably false. Because no whistle-level annotations exist for these recordings, the figure shows what the detector produces; it does not measure its accuracy.

### 3.6. Computational cost

Running the 5–24 kHz detector on a 65-s recording took about 2.8 s on a CPU, about 23 times faster than real time, and running it on all 12 DCLDE 2011 recordings (43.2 min), including reading the files, took about 4 min on a free Colab CPU. Silbido processed the 12 recordings in about 90 s in MATLAB on a different computer, so these times are not a direct speed comparison.

## 4. Discussion

### 4.1. Recall and precision

Trained only on synthetic whistles in noise from a different ocean region, species and recording system, the network transferred well enough to find more of the annotated DCLDE 2011 whistles than Silbido, on 9 of the 12 recordings, with similar frequency accuracy and somewhat higher coverage. Its precision varied widely between recordings, however (18% to 89%), so how well it transfers depends on the recording. Its weakness is precision: on average about 44% of its detections in the main scoring did not match a counted whistle, against 24% of Silbido’s. Over the benchmark as a whole, the two effects nearly balance, and with only 12 recordings the F1 difference cannot be distinguished from zero.

The false-alarm analyses suggest that the precision gap has more than one cause, and that only part of it reflects sounds that are not whistles. First, a substantial share of RidgeTrace’s sampled false alarms were whistle-related (36 of 80, or 45%): whistles absent from the annotations, or harmonics, pieces and echoes of annotated whistles (Figure 4b,c). The DCLDE 2011 analysts annotated strong whistles and skipped many very short ones [28], so the annotations are incomplete for short and faint whistles, especially in busy recordings; in `070925-023000` only 7% of the annotated whistles met the scoring criteria at all. Second, the 40-fold difference between RidgeTrace’s false-alarm rates on dolphin recordings and on whistle-free recordings is consistent with most of RidgeTrace’s false alarms occurring when whistles are present, rather than on clicks or ambient noise. The two sets of recordings differ in more than the presence of whistles, however: the whistle-free recordings come from the same survey and equipment as the training noise, and the DCLDE 2011 recordings do not, so part of the difference may reflect that. Third, part of the whistle-related false alarms are real errors of the tracker: breaking one whistle into several pieces, following a harmonic separately, or jumping between whistles at crossings (Figure 1c, near 9.9 s). These are scored as false alarms whatever the annotations’ completeness, and they suggest that the tracker, rather than the network, is the first thing to improve. We did not test this directly, for example by comparing cell-level and track-level performance on annotated recordings or by replacing the tracker, so it remains a hypothesis.

We do not use these observations to adjust the scores. The blind sorting was done by one person on a sample, 63 of the 147 decisions (43%) were marked unsure, and an unannotated whistle can still be an unwanted detection if the goal is to reproduce an analyst’s choices. Silbido’s false alarms would include unannotated whistles too, though fewer of them in the sample.

### 4.2. False alarms without whistles

Where no whistling species was logged, RidgeTrace produced about one false alarm every minute, about five times fewer than Silbido at its default setting. This difference depends on applying the 150 ms minimum to RidgeTrace: before the common filter, the rates returned under each method’s own settings were similar (4.46 vs 5.06 per minute), and Silbido’s already included its built-in 150 ms minimum (Section 3.3). Of the three Silbido thresholds tested, only 12 dB gave a comparably low rate, at which point it found 57% of the DCLDE 2011 whistles instead of RidgeTrace’s 87%. This combination, high recall where whistles are present and few detections where they are not, is useful in practice: in long-term monitoring most recording time contains no whistles, so the false-alarm rate in empty recordings may largely set the cost of reviewing detections. Where dolphins are present, however, RidgeTrace’s review cost is higher than Silbido’s (35.6 vs 10.3 false alarms per minute on DCLDE 2011). It is consistent with the training design: the network was shown loud synthetic clicks and sweeps that were never targets, and the tracker discards broadband frames and pulse-train patterns.

Two caveats apply. The whistle-free recordings come from the same survey and array as the training noise, so RidgeTrace may have an advantage over Silbido there that would not hold in other recordings. And whistle-like sounds have occasionally been reported near beaked whales [53, 54], and distant dolphins could be audible in recordings where none were logged, so a few “false alarms” could be real whistles; the blind check found two possible whistles among RidgeTrace’s 20 sampled detections and none among Silbido’s.

### 4.3. Comparison with other work

RidgeTrace’s approach resembles that of Li et al. [26, 27], who also trained networks on synthetic or automatically generated training data and evaluated on DCLDE 2011. Their best model reported F1 87.20 over 5–50 kHz, well above RidgeTrace’s 67.1 over 5–24 kHz, but the band, the training data and the network all differ, and our Silbido scores differ from theirs in precision, so the numbers cannot be compared directly. Our contribution is not a better score but a set of observations about a very small, cheaply trained model: it appears to transfer across species and recording systems, though with precision that varies widely between recordings; its errors are consistent with being concentrated where whistles are present; and, after applying the 150 ms minimum, it appeared to produce relatively few false alarms in whistle-free recordings from the same survey as its training noise. Evaluating a 5–50 kHz version of RidgeTrace with the same scorer would allow a direct comparison and is planned.

### 4.4. Limitations

- **Settings chosen after seeing test results.** Keeping harmonics and the 150 ms minimum detection length were decided after the first test results. Both have a clear rationale independent of the results (the annotations include harmonics; 150 ms is Silbido’s own default and the annotation criterion), and we report the scores without each (Table 5). No other setting was changed after testing, and nothing was trained or tuned on the test recordings.
- **Frequency band.** The 5–24 kHz band excludes 18% of otherwise-counted whistles, and with all frequencies counted RidgeTrace’s recall advantage (+4.1 points, range *−*6.6 to +13.3) is no longer distinguishable from zero.
- **No operating curve for RidgeTrace.** Silbido was run at three thresholds, but RidgeTrace at only one setting (*τ* = 0.4, minimum mean track probability 0.55), because we chose not to vary its settings on the test recordings. Whether a stricter setting would close the precision gap, and at what cost in recall, is therefore unknown; it should be measured on annotated recordings outside the test set.
- **No comparison with the strongest published method.** Our baseline is Silbido, the established rule-based reference, not the best recent deep-learning contour extractor. We did not run the networks of Li et al. [26, 27] on our test setup, so how RidgeTrace compares with them, over the same band and with the same scorer, is unknown.
- **Small test set.** Twelve recordings, with as few as 9 counted whistles in one (Qx-Dc-SC03), make the bootstrap ranges wide and approximate. After adjusting for three tests, only the recall difference remained below *p* = 0.05.
- **Scoring ambiguity.** The 30% coverage rule is stated by Li et al. [27] but not present in Silbido’s scoring code, and our Silbido precision is lower than published for two species for reasons we could not find.
- **Incomplete annotations.** Some scored false alarms are probably real whistles; we did not change any score because of this.
- **False-alarm sorting.** One person sorted all pictures, only 84 of the 147 DCLDE decisions (57%) were marked sure, and the samples are not proportional to all false alarms.
- **Training data.** All training noise came from one encounter, in which clean windows were scarce; the synthetic contours follow simple rules and may not cover all real whistle shapes. About 20% of the 5–24 kHz synthetic whistles were placed at the 5.4 kHz lower limit because of a fixed starting-frequency range in the generator. The saved checkpoint is from step 2,000 of 6,000, and the minimum mean track probability (0.55, as in the 2–24 kHz detector) was not lowered when the peak threshold was set to 0.4 by the calibration rule on synthetic validation data, before any test run (Section 2.5). We left these as they were rather than change the detector after testing.
- **The 2–24 kHz detector.** The version used to build the dataset for our reproduction of Park et al. [31] has no measured accuracy on annotated real recordings, and its training species (false killer whales) is not among the DCLDE 2011 species. Its synthetic recall and whistle-free false-alarm rate are similar to those of the 5–24 kHz version, and its output on one 2017 recording (Figure 5) follows several clear whistles, but neither is evidence of its accuracy on real whistles.
- **Whistle-free test.** Same survey and equipment as the training noise (different encounters), so it may favour RidgeTrace, which was trained on that noise, over Silbido, which was not adapted to it; no independent whistle-free test was made. Silbido read 16-bit WAV copies of the audio, while RidgeTrace read the original files. The log of species present may also be incomplete.

### 4.5. Future work

The false-alarm analyses suggest the tracker as the first target: joining pieces of one whistle, handling crossings and keeping harmonics with their fundamentals. More varied clicks, burst pulses and noise in the synthetic data may reduce the remaining false alarms. Any such tuning should use annotated recordings outside the 12 test recordings. Other steps are to score the 2–24 kHz detector on DCLDE 2011, to train and evaluate a 5–50 kHz version for direct comparison with published results, to check a sample of detections on the 2017 recordings by hand to estimate precision on false killer whales, and to compare with Silbido’s deep-learning tracker.

## 5. Conclusions

A 14,209-parameter convolutional network trained only on synthetic whistles in real ocean noise, followed by a rule-based tracker, detected 86.9% of the annotated whistles within 5–24 kHz in the DCLDE 2011 test set without any training on annotated data, 14 points more than Silbido (4 points, not clearly different, when whistles at all frequencies are counted). After discarding detections under 150 ms, it produced about one false alarm per minute on recordings with no whistling species present, five times fewer than Silbido; these recordings came from the same survey as the training noise, which may favour RidgeTrace. Its precision on whistle-rich recordings was 20 points lower than Silbido’s, and its overall F1 was 7.5 points lower, a difference that this small test set cannot resolve. In a blind sample from the four recordings where either method performed weakest, 45% of RidgeTrace’s sampled false alarms were unannotated whistles or harmonics, fragments or echoes of annotated whistles, rather than noise, and the tracker appears to be the most promising place to improve. Because it needs no annotated training data and little computing power, the approach may be useful where annotations are lacking, provided its detections are reviewed.

## Acknowledgements

The authors thank the Department of Computer Science and Engineering, Neil Gogte Institute of Technology, Hyderabad, for supporting this research work and for providing an encouraging academic environment. We also thank the organisers of the DCLDE 2011 and DCLDE 2022 workshops and NOAA for making the recordings and annotations publicly available, and the developers of Silbido for making the software and its scoring code available.

## Data availability statement

The DCLDE 2011 recordings and annotations are available from NOAA at https://doi.org/10.25921/wsnc-js16 [28], and the DCLDE 2022 (HICEAS 2017) recordings from NOAA’s public cloud storage [32]. The code, trained models, detections, score files, false-alarm sorting sheets and the scripts that produce every table and figure are available from the corresponding author on reasonable request.

## Ethics statement

This study used only archival recordings made by others; no animals were handled or disturbed for this study.

## Use of generative AI

The study design, data selection, detector development, experiments, Silbido runs and the blind sorting of false alarms were carried out by the authors. Claude (Anthropic; model Claude Opus 5.5) was used to draft and edit the manuscript text and its LaTeX formatting from the authors’ results and notes. It was used to speed up the writing, formatting and checking of the manuscript. Every number reported was taken from the authors’ own result files and checked against them, and the authors reviewed and revised the full text. The authors take full responsibility for the content of this article.

## Appendix A. DCLDE 2011 test recordings

**Table A1.** The 12 DCLDE 2011 recordings of the Li et al. [27] test set, with the short names used in Table 4.

| Species | Short name | Recording |
|---|---|---|
| Bottlenose dolphin | SCI0608 | Qx-Tt-SCI0608-N1-060814-121518 |
|  | 070924-205305 | palmyra092007FS192-070924-205305 |
|  | 070924-205730 | palmyra092007FS192-070924-205730 |
| Long-beaked common dolphin | CC0411 | Qx-Dc-CC0411-TAT11-CH2-041114-154040-s |
|  | FLIP0610 | QX-Dc-FLIP0610-VLA-061015-165000 |
|  | SC03 | Qx-Dc-SC03-TAT09-060516-171606 |
| Melon-headed whale | 071004-032342 | palmyra092007FS192-071004-032342 |
|  | 061020-204327_4 | palmyra102006-061020-204327_4 |
|  | 070925-023000 | palmyra092007FS192-070925-023000 |
| Spinner dolphin | 070927-224737 | palmyra092007FS192-070927-224737 |
|  | 061103-213127_4 | palmyra102006-061103-213127_4 |
|  | 071011-232000 | palmyra092007FS192-071011-232000 |

## Appendix B. Reproducibility

All scores in this paper were reproduced on a fresh machine from the stored detections: the 12 recordings and both annotation sets matched their published sizes and checksums, the stored detection files were byte-identical, the scorer’s self-test gave the expected result, and all 11 score files were reproduced field by field. Every table and figure is produced by a script from the stored result files. Software: Python 3.11 and 3.13, PyTorch 2.11, SciPy; Silbido 3.0 in MATLAB R2024b.

